# Development of a Novel Methyl Cellulose Hydrogel with Physiologically-Relevant Controlled Ethanol Release for Cervical Dysplasia Ablation

**DOI:** 10.1101/2025.11.25.690497

**Authors:** Ashleigh M Jankowski, Erela Imanoel, Gatha Adhikari, Jenna Mueller, Katharina Maisel

**Affiliations:** Fishell Department of Bioengineering, University of Maryland, College Park; Department of Obstetrics, Gynecology & Reproductive Science, University of Maryland School of Medicine, Baltimore, MD, USA; Marlene and Stewart Greenebaum Cancer Center, University of Maryland School of Medicine, Baltimore, MD, USA

**Keywords:** topical gel, cervical dysplasia, ethanol ablation

## Abstract

Cervical cancer is a leading cause of death in women in low- and middle-income countries (LMICs) and disproportionately affects women of minority populations in the US, primarily due to a lack of infrastructural support for specialized care. A promising treatment which meets accessibility requirements is ethyl cellulose (EC)-ethanol ablation—inducing necrotic cell death through application of ethanol to kill precancerous cells. While previous work focused on injecting EC-ethanol to ablate high-grade dysplasia (which can reach depths up to 5 mm below the tissue surface), low-grade dysplasia requires a different delivery method as it is much more superficial (reaching depths of only 1-3 mm). Here, we have developed a topical gel for local ethanol ablation of low-grade dysplasia with minimal damage to healthy cervical tissue. We investigated several gellants, including methyl cellulose (MC), EC, and Pluronic® F-127, to develop an ethanol gel that meets parameters for low cost and topical ease of use. Formulations with F-127 did not form gels with ethanol. Formulations with EC and MC were gel-forming. The MC-based formulations formed more uniform and stable gels that hold their own weight while still being spreadable at both room and body temperatures, key criterion for local cervical application. The optimal formulation contained 70% ethanol, 20% water, and 10% MC. One gram of this formulation represents approximately 5¢ material cost, and formulated gels were stable for one week at least when stored at 4, 22, 30, and 37 °C. Additionally, the MC gel achieved localized ablation within 5 minutes after application to cervical cancer cells *in-vitro*. Taken together, we have developed a low-cost, efficacious, MC-based ethanol gel fit for translational testing to treat low-grade cervical dysplasia. This gel may provide a novel treatment option for women in LMICs, without causing major side effects or loss of healthy cervical tissue.

**Translational Impact Statement:** We have developed a low-cost, efficacious, methyl cellulose-based ethanol gel fit for translational testing to topically treat low-grade cervical dysplasia. This addresses the need for a novel and accessible treatment option for women in low- and middle-income countries, which will not cause major side effects or loss of healthy cervical tissue.

**Ethics Statement:** All work herein are the authors own original work. Authors have no conflicts of interest to disclose. Primary funding for the project was startup funds provided by the University of Maryland. All data available upon request.

## I. Introduction

Cervical cancer incidence is 2-3 times higher and mortality is 3-6 times higher in low- and middle-income countries (LMICs) compared to high-income countries (HICs).^1–4^ Disparities in cervical cancer incidence and mortality also exist within HICs, with significant disparities occurring in populations with poor access to reproductive health services in the United States, most notably rural populations and historically marginalized racial groups.^5–9^ There is a need for better solutions across the cervical cancer care continuum, particularly in increasing access to treatment. For women diagnosed with low-grade dysplasia (i.e. pre-cancer), a ‘wait and see’ approach is typically used in high-income communities to see if lesions resolve on their own; however, in low-income communities, women are often lost in the process of follow up care, which can allow for low-grade dysplasia to develop into later stage cervical pre-cancer or cancer. Because of this, the World Health Organization and others recommend “see-and-treat” approaches to enable cervical cancer prevention— i.e., patients in low-resource settings whose pre-cancer are diagnosed and treated in the same day have better outcomes.^10^

In high-income communities, high-grade dysplasia is commonly treated through surgical excision. While effective, this approach removes substantial amounts of healthy cervical tissue and can lead to adverse obstetric outcomes and other adverse side effects.^11^ A more targeted alternative is ethanol ablation which involves direct application of ethanol to malignant tissue, inducing cell death through necrosis.^12^ Ethanol ablation is particularly well-suited for LMICs due to its low cost, portability, and ease of use.^13^ Promising pilot studies have demonstrated that combining ethanol with the gelling agent ethyl cellulose (EC) enhances local tissue necrosis while minimizing off-target damage.^14^ While previous research has focused on EC-ethanol injection for treating high-grade dysplasia (which can extend up to 5 mm below the epithelial surface), low-grade dysplasia is more superficial, typically confined to 1–3 mm depths, and therefore requires a different delivery strategy.^15^ To address this gap, we are developing a topical formulation of gel-based ethanol ablation optimized for low-grade dysplasia. This formulation would complement existing injectable treatments, enabling a more comprehensive therapeutic approach to cervical disease in LMICs. While several vaginal gel products exist—such as BufferGel and our own hypotonic thermogel, they are primarily water-based and designed for small molecule delivery while preserving vaginal health. The addition of ethanol to these formulations disrupts their gel structure and adhesive properties. To overcome this limitation, we have developed a novel methyl cellulose–based ethanol gel that forms rapidly upon mixing and adheres to surfaces, including wet tissue phantoms. This formulation exhibits a well-defined sol-gel transition and retains its gel-like consistency at body temperature (37 °C). It is also shear-thinning, allowing for easy application using a custom-designed applicator we propose to develop. Importantly, both the methyl cellulose matrix and ethanol therapeutic are low-cost and widely available, making this solution well-aligned with the resource constraints of LMICs. Many gels have been developed for topical vaginal drug delivery for controlled drug release. Typically formulated from water-soluble polymeric materials such as xanthan gum, carboxymethyl-cellulose, hydroxyethyl cellulose, or carbopols, these materials mix with cervicovaginal mucus in the vaginal canal and provide sustained drug delivery for 12-24 hours.^16^ A number of gels have also been designed for treating cervical cancer through vaginal application, such as a thermosensitive polymer (e.g., Pluronic® F-127) gel developed specifically to treat HPV-induced cervical cancer,^17^ nanoparticle-loaded and small-molecule loaded poloxamer 188/407 (F-127/F188)-based temperature-sensitive in situ gels,^18,19^ and a retinyl-acetate delivering gel.^20^ These gels generally deliver expensive drugs to the entire vaginal tissue (not specifically the cervix) and usually require repeated application, making them less suitable for treating low-grade lesions in low-resource settings. Some existing topical cancer treatments have been suggested for the off-label treatment of cervical cancer, notably Imiquimod and 5-fluorouracil creams.^21^ These creams, however, have largely been investigated for late-stage dysplasia treatment, as they trigger intense physiological responses, which would be unduly aggressive for low-grade dysplasia. Furthermore, these gels are water-based and made for delivery of small molecule therapeutics; adding ethanol disrupts their solid-like gel structure and stickiness, making adapting an existing formulation insufficient for ethanol ablation applications. As such, there is a need to develop an ethanol-based gel specifically for cervical application for treatment of low-grade dysplasia.

An ethanol-based gel for topical ablation of low-grade cervical dysplasia will need to have sufficient rheological properties for topical application: the ability to hold its own weight, maintain a storage modulus greater than loss modulus, and high (>100 Pa*s) viscosity maintained at both body temperature (37 °C) and room temperature (22 °C). Further, to meet accessibility requirements, the gel will need to maintain these properties after storage under varying conditions and for extended periods of time, while being low-cost and of simple formulation. Finally, the gel must release ethanol at a rate relevant for achieving kill of cervical dysplasia cells. To develop this gel, we investigated multiple gellants, including Pluronic® F-127, due to its popularity in literature for medical gels; ethyl cellulose (EC), due to its demonstrated capability to form ethanol-releasing gels; and methyl cellulose (MC), due to its demonstrated ability to form ethanol-based gels that maintain properties at varying temperatures, all of which are FDA-approved generally regarded as safe (GRAS) materials,^13,22–27^ and, in the case of cellulose gellants, known to be very low-cost.^28,29^ We investigated storage and loss moduli and viscosity with these various formulations and found that 70% ethanol, 20% water, and 10% MC gel best exemplified goal parameters in that it is low-cost and has sufficient viscoelastic properties to hold its own weight while maintaining spreadability for topical application. We then tested this gel for ethanol release and localized *in-vitro* in cervical cancer cell-killing.

## II. Methods

### i. Materials

Methyl cellulose (MC) viscosity 4000 cP (Thermo Fisher, Geel, Belgium) was received from Thermo Fisher Scientific and used without further purification. Ethyl Cellulose (EC) viscosity 100 cP and Pluronic® F-127 (F127) Powder were acquired from Sigma Aldrich, (Sigma-Aldrich, Co., St. Louis, MO) and used without further purification. 200 Proof Ethanol (EtOH) (200 proof, Pharmco) was the primary solvent used other than Millipore Ultrapure deionized (DI) water.

### ii. Gel Formulation

All gel formulation was performed at room temperature. Gellant (MC, EC, or F127) was weighed out to appropriate mass and slowly added to ethanol at desired concentrations (Table 1) while mixing with stir bar or by hand with spatula. Added EC required more time to dissolve than MC/F127, approximately 1h. After adding the entire measured mass of gellant, gels were further mixed using a handheld tissue homogenizer. Gels were stored in 20 mL volume scintillation vials sealed with parafilm at room temperature to avoid ethanol evaporation. Except for specified stability studies, all gels were used within 1 week of formulation.

**Table 1.** Concentrations of Gellants and Solvents Investigated.

| Material | % Gellant (w/w) | % DI Water | % EtOH |
| --- | --- | --- | --- |
| MC | 6-15% | 10-20% | 70-79% |
| EC | 6-40% | -- | 60-94% |
| F127 | 5-35% | 20-80% | 71-82% |

### iii. Rheological Analysis

Solution viscosity and rheological characterization was performed using an Ares G2 Rheometer (TA Instruments, New Castle, DE) equipped with a 40 mm diameter 1.991° stainless steel cone plate geometry and a solvent trap. Viscosity was evaluated through use of oscillation time sweep flow ramps, wherein shear rate was linearly decreased from 250 s^-1^ to 1 s^-1^ over 60 seconds followed by a reverse wherein the shear rate was linearly increased from 1 s^-1^ to 250 s^-1^ over 60 seconds. Storage and loss moduli were evaluated using an oscillation time sweep, wherein an angular frequency of 10 rad/s constant strain of 1% was applied over a duration of 240 seconds. The angular frequency of 10rad/s and strain of 1% were chosen based on common use for similar gels in literature, as well as published analyses that methylcellulose and related cellulose systems maintain linear viscoelastic behavior for strains <5%.^30–32^ All analyses were performed in triplicate at 22 °C or 37 °C. Any samples which were stored in a refrigerator for stability study purposes were also evaluated at 4 °C. Formulations were defined as a successful gel if they met specific criteria: a storage modulus (G’) significantly greater than a loss modulus (G’’), time-dependent viscoelastic response under applied shear which showed a yield stress, and qualitatively able to hold their own weight (evaluated via inversion in scintillation vial) while still being spreadable (easily spread into a homogenous even layer).^33–36^

### iv. Release Studies

Release studies were performed to evaluate the ethanol delivery capabilities of the developed gels. Gels were loaded into 10 mm flat width dialysis tubing cellulose membrane (Sigma Aldrich, St. Louis, MO) sealed with dialysis tubing clips (Spectrum™ Dialysis Tubing Closures: Spectra/Por™ Standard 12 mm), and submerged in 1X PBS at 37 °C in a sealed container on a tube rotator (50 mL centrifuge tubes filled to reduce air volume, final PBS volume of 53 mL). 3x 300ul samples were taken at set time points over the course of 6 hours. Ethanol concentration was detected using an Amplite® Ethanol Quantitation Kit (AAT Bioquest, Pleas-anton, CA) and a Spark Multimode Microplate Reader.

### v. Stability Studies

Stability studies were performed by storing gels at various temperatures and evaluating the changes in rheology over time. Temperatures included 4 °C, as the standard refrigeration temperature, 22 °C, as average room temperature, 30 °C, as the highest recorded average room temperature in sub-Saharan African LMICs, and 37 °C, to simulate high temperature outdoor conditions in sub-Saharan African LMICs. ^37^ These studies were designed to simulate real-world storage and transport conditions, taking into account that resource-poor settings often have limited access to climate-controlled storage. 10 mL of gel were prepared and stored in 20 mL scintillation vials, which were sealed. Gel mass and rheological characterization were recorded at set intervals - mass every day for a week and rheological characterization at completion of the trial period. The trial period was performed for 1 week in order to confirm that gels maintained stability from formulation time to use over the course of studies, as well as to begin validation for storage in a clinical setting (long-term studies currently ongoing). Rheological characterization was performed at 22 °C as described above. Gels were also qualitatively analyzed to assess if clumping or ethanol-gellant separation occurred. Parameters to identify a gel as stable were viscosity peak, which is not significantly different from the identified average peak for that gel when made fresh, a storage modulus which is greater than loss modulus, no statistically significant changes in mass, and no observed gel separation.

### iv. *In-Vitro* Cell Killing Assay

A human cervical cancer cell line, SiHa, from ATCC (ATCC® HTB-35™, Manassas, VA) was used to determine ethanol killing. SiHa cells were expanded in Eagle’s Minimum Essential Medium (EMEM, ATCC), supplemented with 10% FBS (Sigma-Aldrich, St. Louis, MO) and 1% Penicillin-Streptomycin (P/S, Sigma-Aldrich). Cells were expanded in standard cell culture conditions (37 °C, 21% O2, 5% CO2) in 6 well plates. Cells were recovered at 80% confluency. 10 µL of the gel, 1X phosphate-buffered saline (PBS) or 200 proof ethanol were added to the designated wells. Cells were exposed to gels/control for 5 minutes. This timescale was chosen as it is more than enough time for ethanol killing of a single cell layer,^38^ while still being short enough that cells can survive without addition of media, a potential vector for ethanol spread. Viability was then assessed with a live/dead assay by adding calcein AM (2M, eBioscience™ Calcein AM Viability Dye, 65-0853-39, San Diego, CA) and propidium iodide (5M Propidium Iodide Ready Flow™ Reagent, R37169) to the cells for 15 minutes. Cells were imaged at 10X using a Nikon Ti2 fluorescence microscope. Next, live and dead cells in each treatment group were counted in ImageJ Fiji (NIH, Bethesda MD). For each image live and dead channels were split, then the image was thresholded, and a binary image was obtained. Then the analyze particle tool was used and cells were counted with a manually created ROI. All studies were performed in triplicate.

## Results

### F127 does not form gels in combination with ethanol

F127 as a gellant for ethanol-based gels was evaluated across a range of concentrations [5-35% (w/w)]. Concentrations lower than 15% did not form gel-like materials at room temperature, which aligns with literature understanding of F127 critical gelling concentrations,^39^ and concentrations higher than 25% tended to approach more rubbery, solid-like textures. F127 combined with 200 proof ethanol alone did not form a homogenous mixture; the components stayed separated. Gels which successfully formed homogenous formulations were evaluated for viscosity and storage/loss moduli. For all formulations and temperatures, viscosity peaks were less than 10 Pa*s, indicating that the formulations were not forming a gel: the maximum viscosity achieved in independent F127 gels was the 10% F127, 20% water, 70% ethanol formulation at 0.054 +/- 0.007 Pa*s. The 15% F127, 20% water, 65% ethanol formulation peaked at 2.7 Pa*s; 15% F127, 85% ethanol peaked at 1.9 Pa*s (**Figure 1A-C**). These peaks also occurred at the beginning of a viscosity shear sweep, after which the viscosity drastically decreased for all, indicating shear-thinning behavior. For all evaluated formulations and temperatures containing ethanol (15% F127, 20% water, 65% ethanol; 15% F127, 85% ethanol; 10% F127, 20% water, 70% ethanol; at 37 °C and 22 °C), the loss modulus was greater than the storage modulus, indicating the formulations behaved more like a viscoelastic liquid than a gel (**Figure 1D-I**). Additionally, the moduli did not stay the same when at both 22 °C and 37 °C for all formulations, indicating a temperature-dependent viscoelastic profile. The only F127 formulation which formed a successful gel was that which did not include ethanol, the 20% F127 80% water formulation. This formulation displayed viscosity above 3900 Pa*s for all temperatures and a storage modulus greater than loss modulus for all temperatures (**Figure S1A-C**). Overall, F127 gels lacked the properties necessary for sufficient gelation: ability to hold their own weight, time-dependent viscoelastic response under applied shear which showed a yield stress, and inability to maintain gel properties at room and body temperatures.

**Figure 1.**
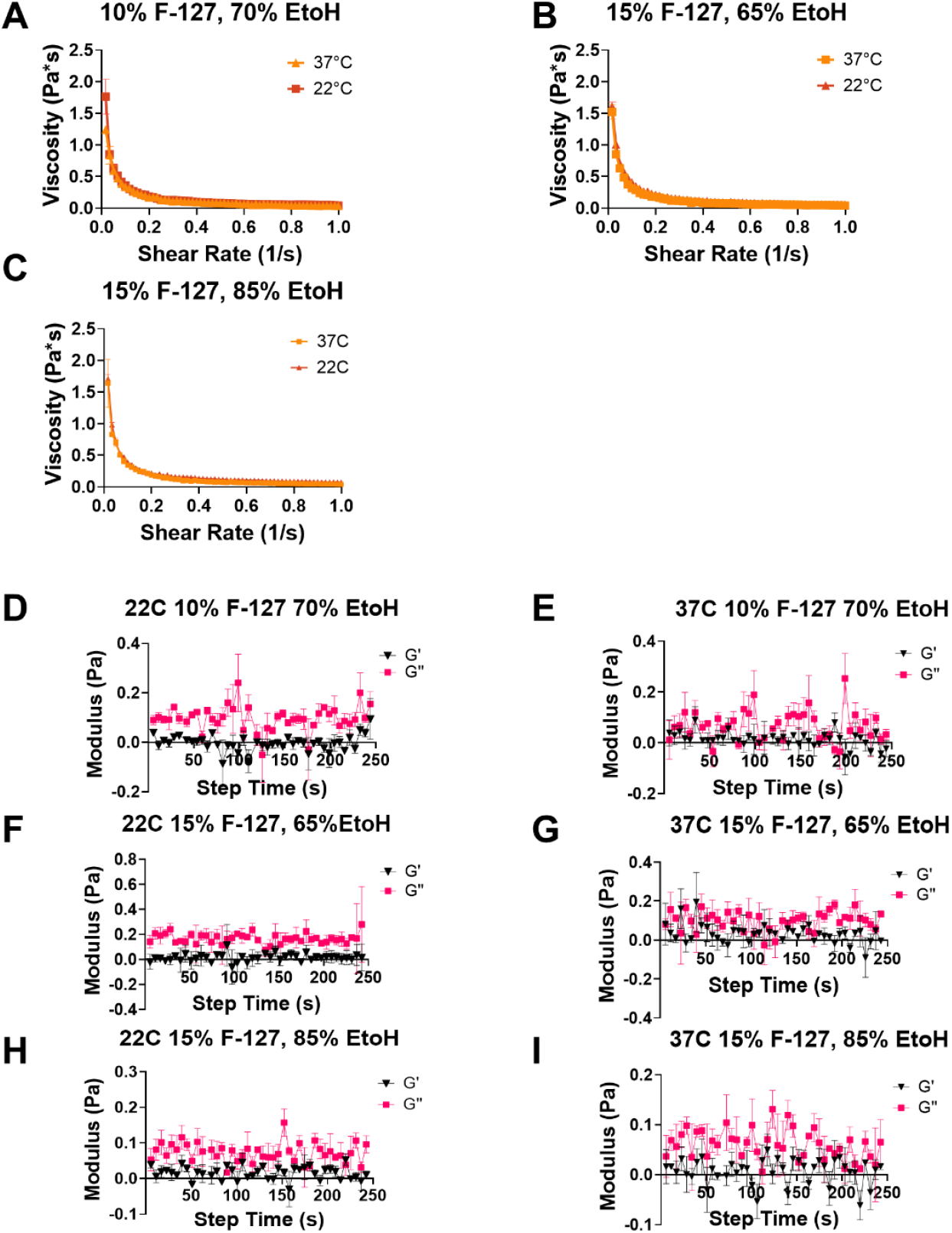
Rheological properties of Pluronic® F-127 Gel formulations. Average ± SEM values for viscosity-stress curves (top) and storage-loss moduli (bottom). Data shown for formulations of 10% Pluronic® F-127, 20% water, 70% ethanol (A, D-E), 15% Pluronic® F-127, 20% water, 65% ethanol (B, F-G), 15% Pluronic® F-127 and 85% ethanol (C, H-I). N=3 for all formulations.

### High percentage ethyl cellulose formulations formed gels and exhibited temperature-dependent properties

Ethyl cellulose (EC) formulations were made with 6, 10, 15, 20, 25, 30, and 40% EC and ethanol. The viscosity of these formulations greatly decreased when brought up to 37 °C, with little to no peaks, suggesting these formulations lost their gel properties at high temperatures (i.e., they were melting) (**Figure 2A-F**). At EC percentages less than 30%, the loss modulus was greater than the storage modulus at both 22 °C and 37 °C, indicating the formulations were a viscoelastic liquid as opposed to a gel (**Figure 2G-L**). The 40% EC formulation was a non-spreadable near-solid at room temperature. 40% EC data not shown, as formulations were so stiff they were a malleable solid instead of a gel, making rheological testing in the same manner as was done for 6-40% gels nonviable.

**Figure 2.**
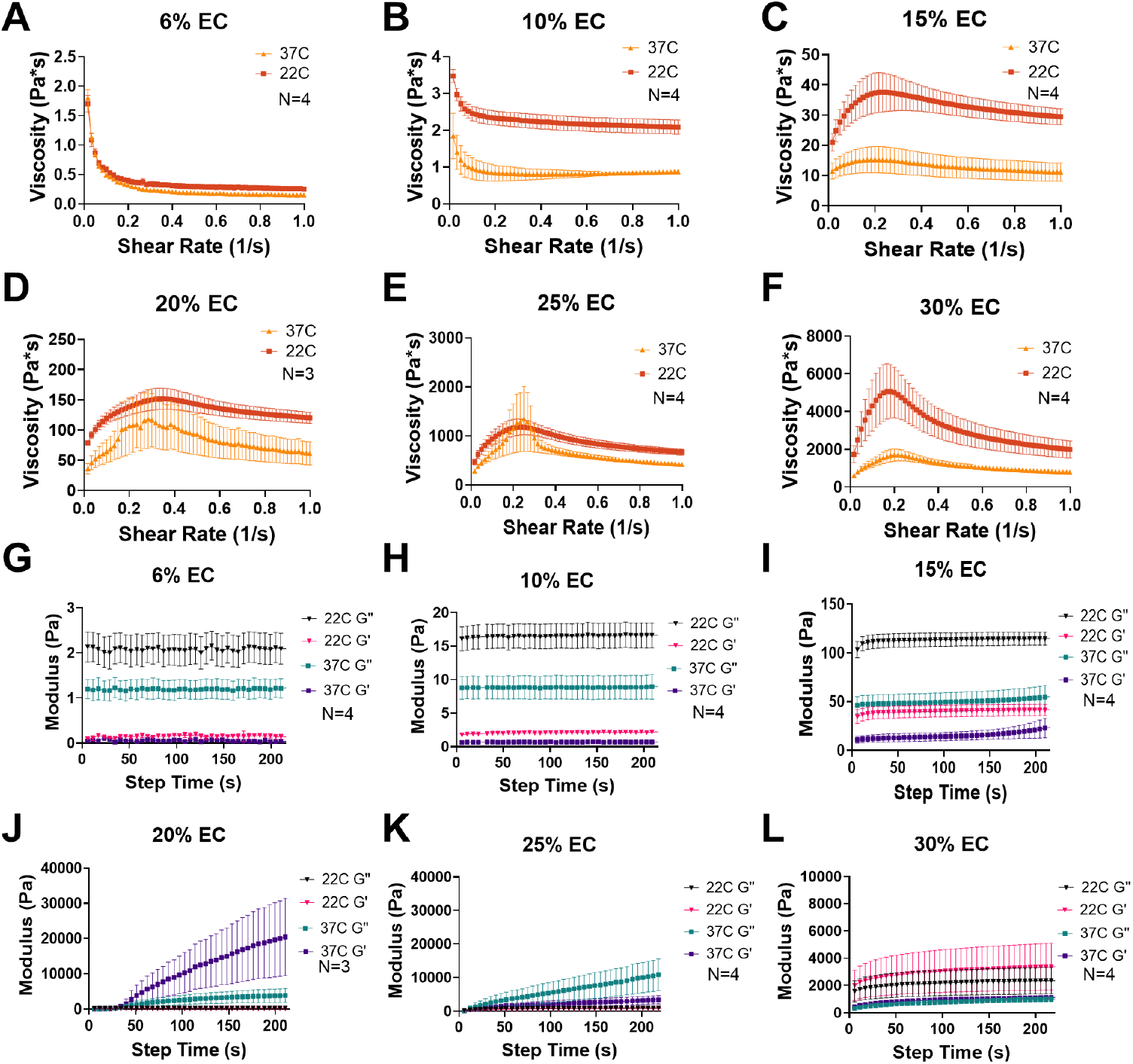
Rheological properties of ethyl cellulose gel formulations from 6-30% EC. Average ± SEM values for viscosity-stress curves at 22 °C and 37 °C (A-F), and storage and loss modulus oscillatory time sweeps (G-L).

### Methylcellulose formulations formed temperature-stable high-viscosity gels

Methyl cellulose (MC) formulations were tested at a broad range of MC, water, and ethanol concentrations to identify the maximum and minimum concentrations that could be used to formulate a homogenous solution. Mixtures with >10% MC and/or < 20% water had incomplete dissolution of MC and did not form a homogenous gel. Mixtures with >75% ethanol tended to separate. The MC formulations within these ranges at all concentrations had storage moduli (G’) significantly greater than loss moduli (G’’) (**Table 2**). However, only certain formulations exhibited a time-dependent viscoelastic response under applied shear which showed a yield stress (**Table 2**). Formulations which exhibited a yield stress and had G’>G” were considered to have formed a viable gel.

**Table 2.** Methyl cellulose Formulations Rheological Data.

| % MC | % Water | % EtOH | Exhibited Yield Stress? | $G' > G''$ at 37°C? | Maximum Viscosity at 22 °C (Pa*s) | Maximum Viscosity at 37 °C (Pa*s) | Formed Gel? |
| --- | --- | --- | --- | --- | --- | --- | --- |
| 6 | 20 | 74 | No | Yes | 5550 | 15240 | No |
| 6.5 | 15.5 | 76 | No | Yes | 2698 | 17670 | No |
| 8 | 13 | 79 | Yes | Yes | 15480 | 73240 | Yes |
| 9 | 14 | 77 | No | Yes | 18140 | 18250 | No |
|  | 17 | 74 | No | Yes | 8807 | 10550 | No |
|  | 14 | 77 | No | Yes | 18140 | 18250 | No |
| 9.5 | 14.5 | 76 | No | Yes | 38690 | 32080 | No |
|  | 18.5 | 72 | No | No | 7884 | 14700 | No |
|  | 14.5 | 76 | No | Yes | 38690 | 32080 | No |
|  | 18.5 | 72 | No | Yes | 7884 | 14710 | No |
|  | 14 | 76.5 | Yes | Yes | 51220 | 27720 | Yes |
|  | 15 | 75.5 | No | No | 788.7 | 590.2 | No |
|  | 14.5 | 76 | No | Yes | 38690 | 32080 | No |
| 10 | 20 | 70 | Yes | Yes | 2558 | 2265 | Yes |
|  | 14.5 | 75.5 | Yes | Yes | 526.9 | 3013 | Yes |
|  | 15 | 75 | Yes | Yes | 1129 | 2157 | Yes |
| 15 | 10 | 75 | Yes | Yes | 5600 | 981.3 | Yes |

The formulation with the most consistency in measurements and that qualitatively had texture best able to be spread into an even layer while still holding its own weight was the 10% MC, 20% water, 70% ethanol formulation. This gel had viscosity peaks between 2000-2500 Pa*s at both 22 °C and 37 °C (**Figure 3A**). A peak indicating yield stress is exhibited at 74.9 Pa, and post-yield stress as the shear rate increases, viscosity declines and stabilizes between 200-270 Pa*s (**Figure 3A**). Across both 22 °C and 37 °C, G′ consistently exceeded G′′ by a factor of >3-4, with no significant difference between rheological measurements (**Figure 3B-C**) and the gel formed a homogenous mixture that holds its own weight (**Figure 3D**).

**Figure 3.**
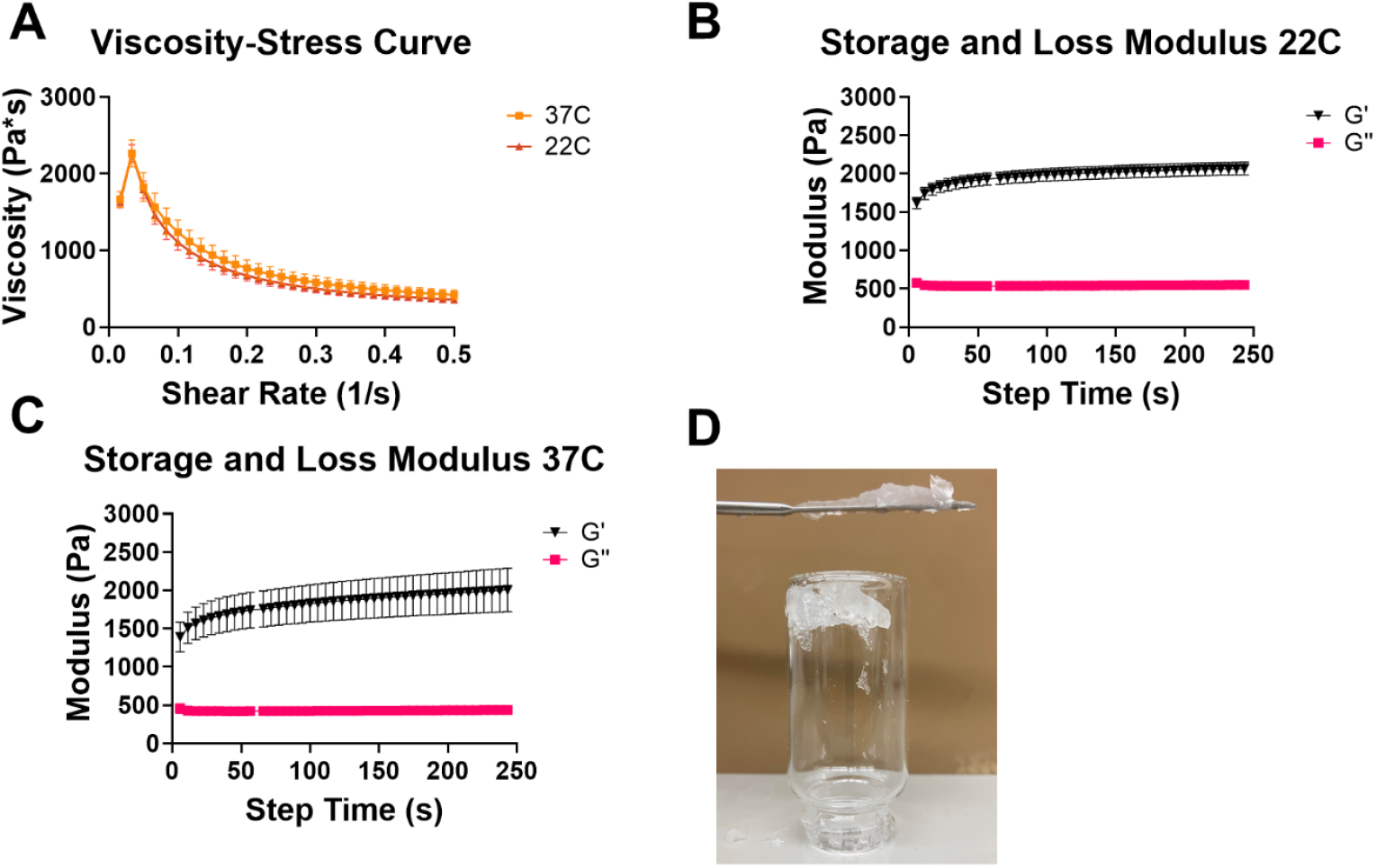
Average rheological data for 20% MC, 10% water, 70% ethanol formulation. Average ± SEM values for viscosity-stress curve (A) and storage and loss modulus over oscillatory time sweep at (B) 22 °C and (C) 37 °C. N=10. (D) Representative image of gel holding its own weight on spatula and in inverted scintillation vial.

### 10% MC gels release 70% ethanol content after 3h and gels are stable for at least one week

Ethanol release for the formulation with optimal rheological properties (10% MC, 20% water, 70% ethanol) occurred in < 4 hours, after which equilibrium was reached (**Figure 4**). Approximately 72 ± 16% of the loaded ethanol was released, indicating high efficacy for controlled ethanol release. A dip in percentage released was observed around 5h., likely due to evaporation of ethanol from the sink after repeated opening and resealing of the container.

**Figure 4.**
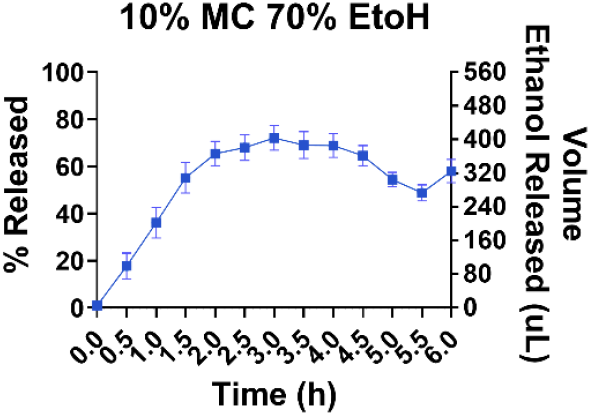
Release over time of ethanol from 10% MC, 20% water, and 70% ethanol gel into PBS. Average ± SEM values for release of ethanol from 1130 mg of 10% MC, 20% water, and 70% ethanol gel formulation in PBS at 37 °C. N=3.

The formulation maintained gel properties over the course of one week. When stored at room temperature for 7 days, yield stress viscosity peak was maintained above 2000 Pa*s (**Figure 5A-B**) and storage modulus was still greater than loss modulus when evaluated at both 22 °C and 37 °C on the seventh day (**Figure 5C-D**). Samples taken throughout the week on days 3 and 5 to verify consistency showed similar trends (**Figure S2A-D**).

**Figure 5.**
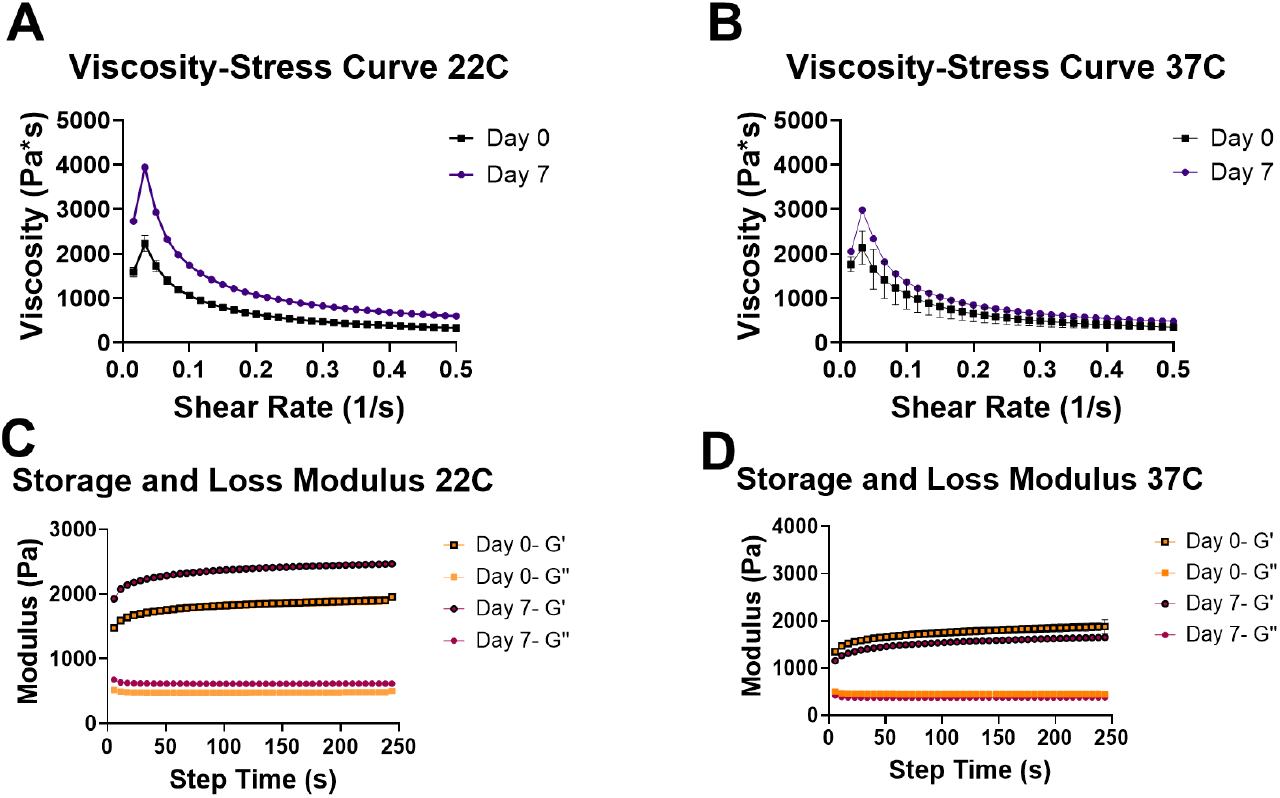
One-week stability study rheology data. Average ± SEM values viscosity stress curves at 22 °C (A) and 37 °C (B) and storage and loss moduli at 22 °C (C) and 37 °C (D) for 10% MC, 20% water, 70% ethanol gel stored at room temperature on days 0 and 7. N=3.

### MC-ethanol gel locally kills SiHa cells within five minutes of gel exposure

The developed gel was then evaluated for cell killing abilities on a human cervical squamous cell carcinoma cell (SiHa) monolayer. Images of SiHa monolayers revealed differential cell viability outcomes based on the applied solution (**Figure 6A**). The PBS control showed no observable cell death (**Figure 6A**). The ethanol-only control (70% ethanol without MC) led to complete cell death across the treated area (**Figure 6B**). MC gel (10% MC, 20% water, 70% ethanol) resulted in an average of 388 dead cells out of 2,722 total and a total cell death area of 0.785 cm^2^ out of 3.06 cm^2^ total area. (**Figure 6C-D**) shows cell death in the area where the gel was applied while leaving the surrounding cells alive. With MC gel cell kill was observed in 25% of the total area whereas the PBS control had 0% cell kill and ethanol had 100% cell kill.

**Figure 6.**
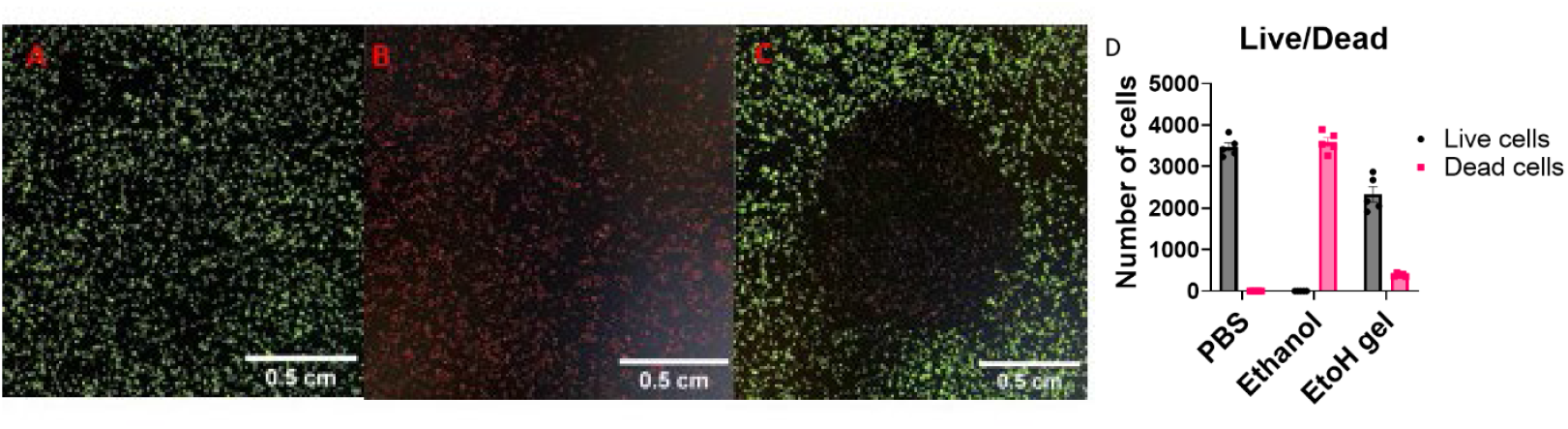
Cell viability outcomes in SiHa monolayers. Microscopy images (A-C) and total cell number (D). Average ± SEM values, N=5

## II. Discussion

Current ethanol ablation techniques for tumors primarily rely on direct ethanol injection into the lesion. While effective for some tumor types, these methods are not suitable for treating low-grade cervical dysplasia, as they pose a significant risk to surrounding healthy tissue.^40^ Previous work successfully developed an injectable ethanol gel for the ablation of high-grade cervical dysplasia, offering a low-cost and targeted treatment capable of inducing sufficient necrosis while preserving adjacent healthy tissue.^22^ However, a holistic cervical cancer prevention strategy for LMICs also requires a safe and effective approach for treating low-grade dysplasia, which is more superficial, typically confined to depths of 1-3 mm.^15^ To address this, we are developing a topical ethanol gel specifically designed to kill tissue only at the site of application, sparing the rest of the cervical surface. This contrasts with current clinical methods such as LEEP, cryotherapy, and thermocoagulation, which often treat the entire cervical face, leading to overtreatment and increased risk of complications. The previously developed injectable gel is not suitable for topical use, as its rheological properties do not allow it to hold its own weight or adhere well when applied to a surface. In contrast, our newly developed gel formulation possesses the mechanical properties needed to remain in place on wet tissue surfaces, delivering localized ablation without specialized equipment. This innovation holds the potential to significantly improve clinical outcomes for women in LMICs and other resource-limited settings. Of the gellants investigated, only MC was found to form an ethanol-based gel which maintained its properties at room and body temperatures. The F127 and EC gels containing ethanol failed to gel and/or maintain consistent properties at both temperatures, making these gellants nonviable for treating low-grade cervical dysplasia lesions locally. The MC formulation which best exhibited the required properties was the formulation containing 10% MC, 20% water, and 70% ethanol. This finding is consistent with similar gels of slightly varied ethanol and methyl cellulose concentrations which have also demonstrated extreme resistance to temperature change.^25^ At all temperatures, this formulation exhibited high viscosity as well as time-dependent viscoelastic response under applied shear which showed a yield stress, has a storage modulus greater than loss modulus (indicating a viscoelastic solid and thus gel formation), was spreadable, and held its own weight, properties which align with those defined for medical use semisolids,^41^ and these properties were maintained at both room and body temperature. This is unique from most common high-ethanol content (>70%) gels, which are typically used for topical bactericide and virucide and as such are less viscous and do not retain gel properties at all temperatures.^42,43^

Release of ethanol from the gel was evaluated to provide insights into whether the gel will release ethanol at a rate relevant for cell-kill under physiological conditions. The release of 72% of the loaded ethanol over the course of four hours indicates a slow release of the majority of the loaded ethanol, which is successful for controlled, local cell killing. Further, the volume of ethanol released from the gel aligns with the volume of ethanol needed to achieve tumor killing in other models.^13^ The volume of ethanol required for tumor cell kill is proportional to the tumor size, with tumors less than 2.5 cm in size requiring as low as 0.1 mL ethanol to achieve cell kill according to current literature.^44^ Given that the average size of cervical dysplastic regions at diagnosis is 6.2 mm^45^, it can be approximated that less than 115 mg of our topically applied gel would be needed for ablation of these lesions. Other developed gels which aim to address cervical cancer topically via release of small molecule drugs tend to require larger doses (up to 2.5 g) and repeat administration (i.e., weekly for 10 weeks), ^46,47^ which would not meet see-and-treat requirements. Further, 115mg of gel represents less than one-cent worth of material costs, sufficiently surpassing The World Banks’ requirements for cost-effective interventions for LMICs, defined as less than 100 USD per year of life saved.^48^ The gel was additionally evaluated for accessibility requirements through its ability to maintain gel properties after a week-long storage at various temperatures, representative of refrigeration storage and room temperature storage in various climates. The gel maintained its properties in all storage conditions, with little to no loss of ethanol due to evaporation or separation during the storage period.

Our optimal gel formulation was shown to release volumes of ethanol at a controlled rate which will be significant for cell kill. The gel enables controlled cell kill in SiHa monolayers by modulating ethanol diffusion. The application of the MC gel (10% MC, 20% water, 70% ethanol) resulted in partial cell death, with an average of 388 cells affected within an area of 0.785 cm^2^, only in the applied region of interest. In contrast, the ethanol-only control led to complete cell death across the treated area, indicating that ethanol alone rapidly diffuses and causes widespread cytotoxicity. The ability of the gel to control ethanol release into only areas of direct application further verifies its viability as a treatment, as this will significantly limit adverse off-target affects, a critical piece of design criteria for cervical dysplasia treatments.^49^ Existing mechanisms for SiHa ablation have comparable monolayer cell kill as the free ethanol group, including application of DOX-loaded AuNR micelles with NIR irradiation^50^ and MoS2-gold nanoparticle supporting nanoribbons for photothermal ablation.^51^ This area-controlled cell death suggests that the MC gel acts as a vital diffusion barrier, slowing ethanol release and limiting its cytotoxic effects. Additionally, this aligns with current understandings of cellulose-based gels with added ethanol components, in that the addition of ethanol enhances tissue penetration depth while cellulose acts as a controlling factor, aiding in targeted cancer cell killing.^52^ This controlled release could be advantageous in applications where partial cell elimination is required, such as targeted therapy or tissue engineering. Future investigation is needed over a longer time scale and in a more complex model to evaluate the penetration depth capabilities of the gel. By preventing excessive ethanol exposure, the MC gel may help preserve surrounding healthy cells while still achieving desired localized cell kill.

## III. Conclusion

The work described herein outlines the development of a novel topical gel formulation for cervical dysplasia ablation. This gel of 10% MC, 20% water, and 70% ethanol is novel in terms of its formulation, properties, and biomedical applications. It has the ability to hold its own weight and the viscoelastic properties of a gel, making it viable for topical applications, a property which it maintains at both room and body temperature. The gel also releases ethanol at a clinically relevant rate causing significant cancer cell death in low-grade cervical dysplasia, indicating significant promise for tumor-ablative properties. The gel further meets requirements for see-and-treat methodology in resource-poor settings such as LMICs and rural localities—inexpensive, shelf stable at a variety of temperatures, and requiring no specialized equipment for use. Future work needs to be done to investigate the efficacy of the gel on a tumor model *in-vitro* beyond just the monolayer, as well as *in vivo* investigations.

## Citations

(1) World Health Organization. Global Strategy to Accelerate the Elimination of Cervical Cancer as a Public Health Problem; 2020.

(2) Singh, D.; Vignat, J.; Lorenzoni, V.; Eslahi, M.; Ginsburg, O.; Lauby-Secretan, B.; Arbyn, M.; Basu, P.; Bray, F.; Vaccarella, S. Global Estimates of Incidence and Mortality of Cervical Cancer in 2020: A Baseline Analysis of the WHO Global Cervical Cancer Elimination Initiative. Lancet Glob Health 2023, 11 (2), e197–e206. 10.1016/S2214-109X(22)00501-0.

(3) Bray, F.; Ferlay, J.; Soerjomataram, I.; Siegel, R. L.; Torre, L. A.; Jemal, A. Global Cancer Statistics 2018: GLOBOCAN Estimates of Incidence and Mortality Worldwide for 36 Cancers in 185 Countries. CA Cancer J Clin 2018, 68 (6), 394–424. 10.3322/CAAC.21492.

(4) Torre, L. A.; Islami, F.; Siegel, R. L.; Ward, E. M.; Jemal, A. Global Cancer in Women: Burden and Trends. Cancer Epidemiology Biomarkers and Prevention 2017, 26 (4), 444–457. 10.1158/1055-9965.EPI-16-0858/174164/AM/GLOBAL-CANCER-IN-WOMEN-BURDEN-AND-TRENDSGLOBAL.

(5) Cohen, C. M.; Wentzensen, N.; Castle, P. E.; Schiffman, M.; Zuna, R.; Arend, R. C.; Clarke, M. A. Racial and Ethnic Disparities in Cervical Cancer Incidence, Survival, and Mortality by Histologic Subtype. Journal of Clinical Oncology 2022, 41 (5), 1059. 10.1200/JCO.22.01424.

(6) Adler, A.; Biggs, M. A.; Kaller, S.; Schroeder, R.; Ralph, L. Changes in the Frequency and Type of Barriers to Reproductive Health Care Between 2017 and 2021. JAMA Netw Open 2023, 6 (4), e237461. 10.1001/JAMANETWORKOPEN.2023.7461.

(7) Lee, H.; Baeker Bispo, J.; Pal Choudhury, P.; Wiese, D.; Jemal, A.; Islami, F. Factors Contributing to Differences in Cervical Cancer Screening in Rural and Urban Community Health Centers. Cancer 2024, 130 (13), 2315–2324. 10.1002/CNCR.35265.

(8) Bhatia, S.; Landier, W.; Paskett, E. D.; Peters, K. B.; Merrill, J. K.; Phillips, J.; Osarogiagbon, R. U. Rural-Urban Disparities in Cancer Outcomes: Opportunities for Future Research. J Natl Cancer Inst 2022, 114 (7), 940–952. 10.1093/JNCI/DJAC030.

(9) Yu, L.; Sabatino, S. A.; White, M. C. Rural–Urban and Racial/Ethnic Disparities in Invasive Cervical Cancer Incidence in the United States, 2010–2014. Prev Chronic Dis 2019, 16 (6). 10.5888/PCD16.180447.

(10) Allanson, E. R.; Schmeler, K. M. Cervical Cancer Prevention in Low- And Middle-Income Countries. Clin Obstet Gynecol 2021, 64 (3), 501–518. 10.1097/GRF.0000000000000629.

(11) Kyrgiou, M.; Mitra, A.; Arbyn, M.; Stasinou, S. M.; Martin-Hirsch, P.; Bennett, P.; Paraskevaidis, E. Fertility and Early Pregnancy Outcomes after Treatment for Cervical Intraepithelial Neoplasia: Systematic Review and Meta-Analysis. The BMJ 2014, 349, g6192. 10.1136/BMJ.G6192.

(12) Shiina, S.; Sato, K.; Tateishi, R.; Shimizu, M.; Ohama, H.; Hatanaka, T.; Takawa, M.; Nagamatsu, H.; Imai, Y. Percutaneous Ablation for Hepatocellular Carcinoma: Comparison of Various Ablation Techniques and Surgery. Can J Gastroenterol Hepatol 2018, 2018 (1), 4756147. 10.1155/2018/4756147.

(13) Morhard, R.; Nief, C.; Barrero Castedo, C.; Hu, F.; Madonna, M.; Mueller, J. L.; Dewhirst, M. W.; Katz, D. F.; Ramanujam, N. Development of Enhanced Ethanol Ablation as an Alternative to Surgery in Treatment of Superficial Solid Tumors. Scientific Reports 2017 7:1 2017, 7 (1), 1–12. 10.1038/s41598-017-09371-2.

(14) Gimenez-Conti, I. B.; Slaga, T. J. The Hamster Cheek Pouch Carcinogenesis Model. J Cell Biochem 1993, 53 (S17F), 83–90. 10.1002/JCB.240531012;PAGEGROUP:STRING:PUBLICATION.

(15) Abdul-Karim, F.; Fu, Y.; Reagan, J.; Wentz, W. B. Morphometric Study of Intraepithelial Neoplasia of the Uterine Cervix. Obstetrics & Gynecology 1982, 60 (2), 210–214.

(16) Cook, M. T.; Brown, M. B. Polymeric Gels for Intravaginal Drug Delivery. Journal of Controlled Release 2018, 270, 145–157. 10.1016/J.JCONREL.2017.12.004.

(17) Bilensoy, E.; Çırpanlı, Y.; Şen, M.; Doğan, A. L.; Çalış, S. Thermosensitive Mucoadhesive Gel Formulation Loaded with 5-Fu: Cyclodextrin Complex for HPV-Induced Cervical Cancer. J Incl Phenom Macrocycl Chem 2007, 57 (1–4), 363–370. 10.1007/S10847-006-9259-Y/FIGURES/7.

(18) Zhang, S.; Zhang, Y.; Wang, Z.; Guo, T.; Hou, X.; He, Z.; He, Z.; Shen, L.; Feng, N. Temperature-Sensitive Gel-Loaded Composite Nanomedicines for the Treatment of Cervical Cancer by Vaginal Delivery. Int J Pharm 2020, 586, 119616. 10.1016/J.IJPHARM.2020.119616.

(19) Zhou, S.; Zheng, X.; Zheng, C.; Qu, F.; Cai, X.; Xu, J. A Thermosensitive Gel Formulation of an Empirical Traditional Chinese Prescription for Treating Cervical Erosion. Acta Pharm Sin B 2012, 2 (5), 495–501. 10.1016/J.APSB.2012.05.005.

(20) Romney, S. L.; Dwyer, A.; Slagle, S.; Duttagupta, C.; Palan, P. R.; Basu, J.; Calderin, S.; Kadish, A. Chemoprevention of Cervix Cancer: Phase I–II: A Feasibility Study Involving the Topical Vaginal Administration of Retinyl Acetate Gel. Gynecol Oncol 1985, 20 (1), 109–119. 10.1016/0090-8258(85)90131-3.

(21) Major, I.; McConville, C. Vaginal Drug Delivery for the Localised Treatment of Cervical Cancer. Drug Delivery and Translational Research 2017 7:6 2017, 7 (6), 817–828. 10.1007/S13346-017-0395-2.

(22) Quang, T. T.; Yang, J.; Kaluzienski, M. L.; Parrish, A.; Farooqui, A.; Katz, D.; Crouch, B.; Ramanujam, N.; Mueller, J. L. In Vivo Evaluation of Safety and Efficacy of Ethyl Cellulose-Ethanol Tissue Ablation in a Swine Cervix Model. Bioengineering 2023, 10 (11), 1246. 10.3390/BIOENGINEERING10111246/S1.

(23) Cadena, I. A.; Adhikari, G.; Almer, A.; Jenne, M.; Obasi, N.; Soria Zurita, N. F.; Rochefort, W. E.; Mueller, J. L.; Fogg, K. C.; Robinson, J.; Mudigunda, S. V.; Keskin, Z. Development of a 3D in Vitro Human-Sized Model of Cervical Dysplasia to Evaluate the Delivery of Ethyl Cellulose-Ethanol Injection. Frontiers in Biomaterials Science 2024, 3, 1365781. 10.3389/FBIOM.2024.1365781.

(24) Akash, M. S. H.; Rehman, K. Recent Progress in Biomedical Applications of Pluronic (PF127): Pharmaceutical Perspectives. Journal of Controlled Release 2015, 209, 120–138. 10.1016/J.JCONREL.2015.04.032.

(25) Lee, D.; Won, J.; Baek, S. W.; Kim, H. Autoignition Behavior of an Ethanol-Methylcellulose Gel Droplet in a Hot Environment. Energies 2018, Vol. 11, Page 2168 2018, 11 (8), 2168. 10.3390/EN11082168.

(26) Gyles, D. A.; Castro, L. D.; Silva, J. O. C.; Ribeiro-Costa, R. M. A Review of the Designs and Prominent Biomedical Advances of Natural and Synthetic Hydrogel Formulations. Eur Polym J 2017, 88, 373–392. 10.1016/J.EURPOLYMJ.2017.01.027.

(27) Nasatto, P. L.; Pignon, F.; Silveira, J. L. M.; Duarte, M. E. R.; Noseda, M. D.; Rinaudo, M. Methylcellulose, a Cellulose Derivative with Original Physical Properties and Extended Applications. Polymers 2015, Vol. 7, Pages 777-803 2015, 7 (5), 777–803. 10.3390/POLYM7050777.

(28) Zou, P.; Yao, J.; Cui, Y. N.; Zhao, T.; Che, J.; Yang, M.; Li, Z.; Gao, C. Advances in Cellulose-Based Hydrogels for Biomedical Engineering: A Review Summary. Gels 2022, 8 (6), 364. 10.3390/GELS8060364.

(29) Seddiqi, H.; Oliaei, E.; Honarkar, H.; Jin, J.; Geonzon, L. C.; Bacabac, R. G.; Klein-Nulend, J. Cellulose and Its Derivatives: Towards Biomedical Applications. Cellulose 2021 28:4 2021, 28 (4), 1893–1931. 10.1007/S10570-020-03674-W.

(30) Eskens, O.; Villani, G.; Amin, S. Rheological Investigation of Thermoresponsive Alginate-Methylcellulose Gels for Epidermal Growth Factor Formulation. Cosmetics 2021, Vol. 8, Page 3 2020, 8 (1), 3. 10.3390/COSMETICS8010003.

(31) Tialiou, A.; Athab, Z. H.; Woodward, R. T.; Biegler, V.; Keppler, B. K.; Halbus, A. F.; Reithofer, M. R.; Chin, J. M. Fabrication of Graded Porous Structure of Hydroxypropyl Cellulose Hydrogels via Temperature-Induced Phase Separation. Carbohydr Polym 2023, 315, 120984. 10.1016/J.CARBPOL.2023.120984.

(32) Zuidema, J. M.; Rivet, C. J.; Gilbert, R. J.; Morrison, F. A. A Protocol for Rheological Characterization of Hydrogels for Tissue Engineering Strategies. J Biomed Mater Res B Appl Biomater 2014, 102 (5), 1063–1073. 10.1002/JBM.B.33088;SUBPAGE:STRING:FULL.

(33) Almdal, K.; Dyre, J.; Hvidt, S.; Kramer, O. Towards a Phenomenological Definition of the Term ‘Gel.’ Polymer Gels and Networks 1993, 1 (1), 5–17. 10.1016/0966-7822(93)90020-I.

(34) Scherer, G. W. Structure and Properties of Gels. Cem Concr Res 1999, 29 (8), 1149–1157. 10.1016/S0008-8846(99)00003-4.

(35) Malkin, A. Y.; Derkach, S. R.; Kulichikhin, V. G. Rheology of Gels and Yielding Liquids. Gels 2023, Vol. 9, Page 715 2023, 9 (9), 715. 10.3390/GELS9090715.

(36) Malvern Instruments Limited. UNDERSTANDING YIELD STRESS MEASUREMENTS; Worcestershire, UK, 2012. https://www.atascientific.com.au/wp-content/uploads/2017/02/MRK1782-01.pdf (accessed 2025-01-18).

(37) Umeh, S. C.; Gil-Alana, L. A. Trends in Temperatures in Sub-Saharan Africa. Evidence of Global Warming. Journal of African Earth Sciences 2024, 213, 105228. 10.1016/J.JAFREARSCI.2024.105228.

(38) Tapani, E.; Taavitsainen, M.; Lindros, K.; Vehmas, T.; Lehtonen, E. Toxicity of Ethanol in Low Concentrations. Experimental Evaluation in Cell Culture. Acta Radiol 1996, 37 (6), 923–926. 10.1177/02841851960373P296.

(39) Alexandridis, P.; Alan Hatton, T. Poly(Ethylene Oxide)-poly(Propylene Oxide)-poly(Ethylene Oxide) Block Copolymer Surfactants in Aqueous Solutions and at Interfaces: Thermodynamics, Structure, Dynamics, and Modeling. Colloids Surf A Physicochem Eng Asp 1995, 96 (1–2), 1–46. 10.1016/0927-7757(94)03028-X.

(40) Koda, M.; Okamoto, K.; Miyoshi, Y.; Kawasaki, H. Hepatic Vascular and Bile Duct Injury after Ethanol Injection Therapy for Hepatocellular Carcinoma. Gastrointest Radiol 1992, 17 (1), 167– 169. 10.1007/BF01888537.

(41) Buhse, L.; Kolinski, R.; Westenberger, B.; Wokovich, A.; Spencer, J.; Chen, C. W.; Turujman, S.; Gautam-Basak, M.; Kang, G. J.; Kibbe, A.; Heintzelman, B.; Wolfgang, E. Topical Drug Classification. Int J Pharm 2005, 295 (1–2), 101–112. 10.1016/J.IJPHARM.2005.01.032.

(42) Sommatis, S.; Capillo, M. C.; Maccario, C.; Rauso, R.; D’Este, E.; Herrera, M.; Castiglioni, M.; Mocchi, R.; Zerbinati, N. Antimicrobial Efficacy Assessment and Rheological Investigation of Two Different Hand Sanitizers Compared with the Standard Reference WHO Formulation 1. Gels 2023, Vol. 9, Page 108 2023, 9 (2), 108. 10.3390/GELS9020108.

(43) Rodrigues, G. B. C.; Filho, J. G. de O.; Jakelaitis, A.; Silva, E. C. da; Placido, G. R.; Marcionilio, S. M. L. de O.; Bitencourt, R. G. Hand Sanitizer Gel Formulations: Influence of Polymer Type on the Rheological Properties. Research, Society and Development 2024, 13 (4), e6313445531– e6313445531. 10.33448/RSD-V13I4.45531.

(44) Paz-Fumagalli, R.; Li, X.; Smallridge, R. C. Ethanol Ablation of Neck Metastases from Differentiated Thyroid Carcinoma. Semin Intervent Radiol 2019, 36 (5), 381. 10.1055/S-0039-1696651.

(45) Tracht, J. M.; Davis, A. D.; Fasciano, D. N.; Eltoum, I. E. A. Discrepant HPV/Cytology Cotesting Results: Are There Differences between Cytology-Negative versus HPV-Negative Cervical Intraepithelial Neoplasia? Cancer Cytopathol 2017, 125 (10), 795–805. 10.1002/CNCY.21905.

(46) Wang, X.; Liu, S.; Guan, Y.; Ding, J.; Ma, C.; Xie, Z. Vaginal Drug Delivery Approaches for Localized Management of Cervical Cancer. Adv Drug Deliv Rev 2021, 174, 114–126. 10.1016/J.ADDR.2021.04.009.

(47) Kirwan, P.; Naftalin, N. J. Topical 5-Fluorouracil in the Treatment of Vaginal Intraepithelial Neoplasia. BJOG 1985, 92 (3), 287–291. 10.1111/J.1471-0528.1985.TB01096.X.

(48) Laxminarayan, R.; Chow, J.; Shahid-Salles, S. A. Intervention Cost-Effectiveness: Overview of Main Messages. 2006.

(49) Desravines, N.; Miele, K.; Carlson, R.; Chibwesha, C.; Rahangdale, L. Topical Therapies for the Treatment of Cervical Intraepithelial Neoplasia (CIN) 2-3: A Narrative Review. 2020. 10.1016/j.gore.2020.100608.

(50) Parida, S.; Maiti, C.; Rajesh, Y.; Dey, K. K.; Pal, I.; Parekh, A.; Patra, R.; Dhara, D.; Dutta, P. K.; Mandal, M. Gold Nanorod Embedded Reduction Responsive Block Copolymer Micelle-Triggered Drug Delivery Combined with Photothermal Ablation for Targeted Cancer Therapy. Biochimica et Biophysica Acta (BBA) - General Subjects 2017, 1861 (1), 3039–3052. 10.1016/J.BBAGEN.2016.10.004.

(51) Pandey, S.; Bansal, A.; Kumar, V.; Mishra, H.; Agrawal, R. K.; Kashyap, S.; Mukherjee, B.; Saxena, P.; Srivastava, A. Gold Nanoparticles Supported on MoS2 Nanoribbons Matrix as a Biocompatible and Water Dispersible Platform for Enhanced Photothermal Ablation of Cancerous Cells Using Harmless near Infrared Irradiation. 2014.

(52) Slavkova, M.; Tzankov, B.; Popova, T.; Voycheva, C. Gel Formulations for Topical Treatment of Skin Cancer: A Review. Gels 2023, Vol. 9, Page 352 2023, 9 (5), 352. 10.3390/GELS9050352.

