## Supplemental Materials for "Development of a Novel Methyl Cellulose Hydrogel with Physiologically-Relevant Controlled Ethanol Release for Cervical Dysplasia Ablation"

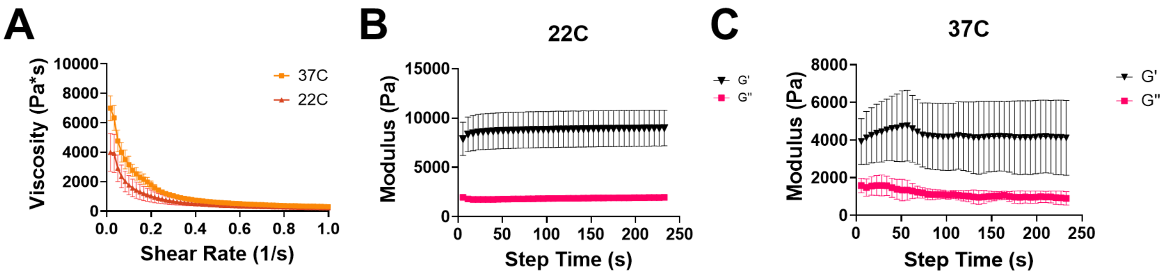


*Figure S1. Rheological properties of 20% Pluronic F-127 80% water formulation. Average ± SEM values for viscosity shear sweep (A) and storage and loss modulus data at 22 (B) and 37C (C). N=3.*

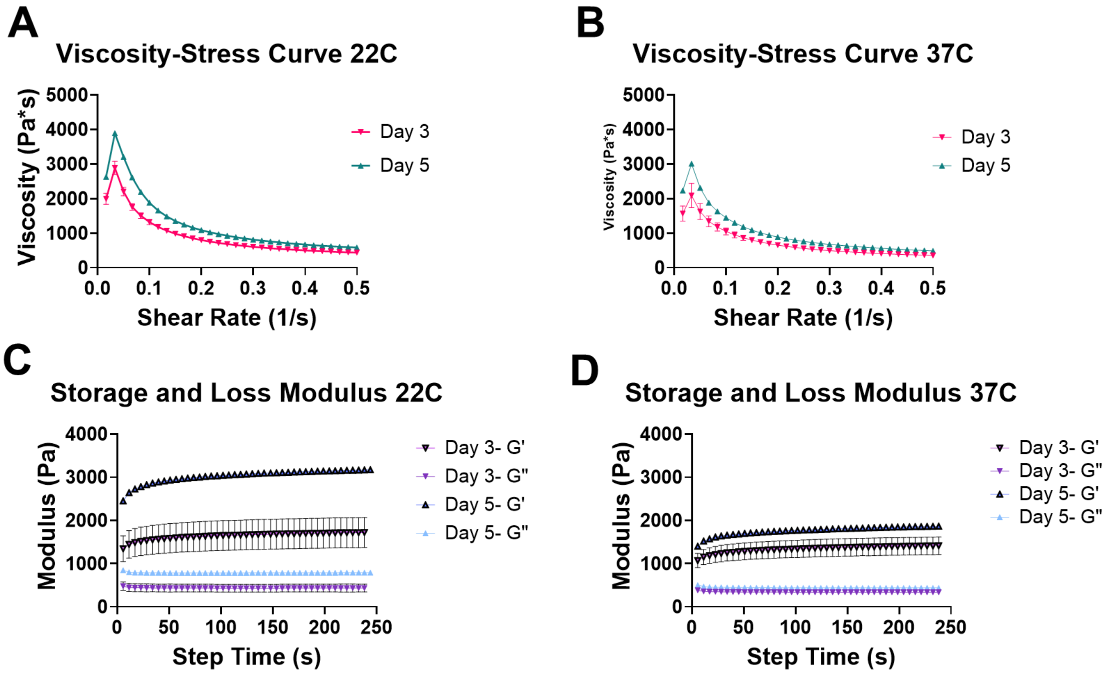


*Figure S2. Day 3 and 5 rheology data from one-week stability study. Average ± SEM values viscosity stress curves at 22C (A) and 37C (B) and storage and loss moduli at 22C (C) and 37C (D) for 70% ethanol 10% MC 20% H2O gel stored at room temperature. N=3.*
